# MetaSRA: normalized sample-specific metadata for the Sequence Read Archive

**DOI:** 10.1101/090506

**Authors:** Matthew N. Bernstein, AnHai Doan, Colin N. Dewey

## Abstract

**Motivation:** The NCBI’s Sequence Read Archive (SRA) promises great biological insight if one could analyze the data in the aggregate; however, the data remain largely underutilized, in part, due to the poor structure of the metadata associated with each sample. The rules governing submissions to the SRA do not dictate a standardized set of terms that should be used to describe the biological samples from which the sequencing data are derived. As a result, the metadata include many synonyms, spelling variants, and references to outside sources of information. Furthermore, manual annotation of the data remains intractable due to the large number of samples in the archive. For these reasons, it has been difficult to perform large-scale analyses that study the relationships between biomolecular processes and phenotype across diverse diseases, tissues, and cell types present in the SRA.

**Results:** We present MetaSRA, a database of normalized SRA sample-specific metadata following a schema inspired by the metadata organization of the ENCODE project. This schema involves mapping samples to terms in biomedical ontologies, labeling each sample with a sample-type category, and extracting real-valued properties. We automated these tasks via a novel computational pipeline.

**Availability:** The MetaSRA database is available at http://deweylab.biostat.wisc.edu/metasra. Software implementing our computational pipeline is available at https://github.com/deweylab/metasra-pipeline.

## 1 Introduction

The NCBI’s Sequence Read Archive (SRA) (Leinonen *et al*., 2011) is a public database that stores raw next generation sequencing reads from over 1.5 million samples belonging to over 75,000 studies. The size and diversity of these samples offers unprecedented opportunity to study the relationships between biomolecular processes and phenotypes across diverse conditions, cell types, and diseases. For example, by focusing on RNA-seq data sets in the SRA, one can study the relationships between gene expression and phenotype. However, such analysis is difficult due to the poor structure of the metadata associated with each sample. The metadata for the biological samples are centrally stored at the BioSample database (Barrett *et al*., 2012). Each sequencing experiment in the SRA references a sample-specific metadata record within this database. The BioSample database organizes each sample’s metadata as sets of key-value pairs where the key is a property of the sample and the value is a property-value of the sample. Figure 1A and Figure 1B display the metadata for two representative samples. Unfortunately, both the properties and property-values are non-standardized and are created at the discretion of the submitter. For this reason, both keys and values consist of synonyms, misspellings, abbreviations, and references to outside sources of information. Furthermore, despite the imposed structure of a key-value description, many of the values are complex, natural language text. For these reasons, aggregate analysis of the data across phenotypes has been difficult. Phenotype-specific studies also remain challenging due to the difficulty in querying for samples that exhibit a phenotype of interest.

**Figure 1:**
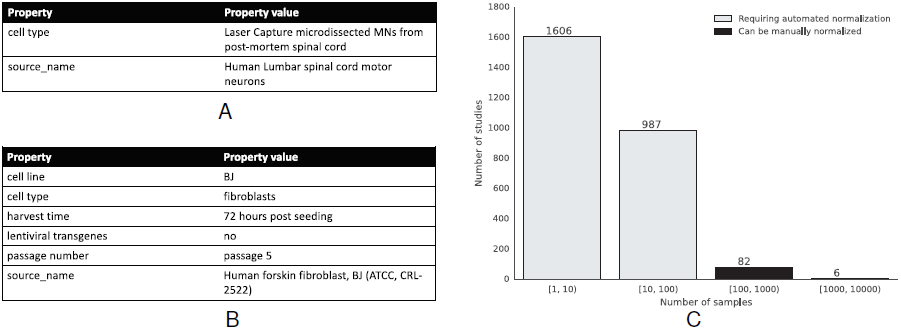
Overview of the dataset. (A) Sample-specific key-value pairs describing sample SRS1217219. Note that the values encode natural language text. (B) Sample-specific key-value pairs describing sample SRS872370. Note the reference to an external cell line BJ. We also note that “forskin fibroblast” is an incorrect spelling. Lastly, the value “no” negates the key “lentiviral transgenes.” (C) Histogram of the number of samples per study for human RNA-seq experiments using the Illumina platform. We assert that the 88 studies each with at least 100 samples can be semi-manually normalized using study-specific methods.

Our goal in this work is to provide structured descriptions of the biological samples used in the SRA. This task is challenging because it requires discriminating between information that describes the biological sample from information that describes other entities such as the sample’s study, sequencing protocol, and lab. A solution to structuring the sample-specific information must address the metadata’s semantics.

Existing methods for normalizing biomedical text focus on annotating the text with terms in a controlled vocabulary, usually in the form of a biomedical ontology. One can approach the task of annotating metadata using either a manual or automated approach. Manual annotation allows for high accuracy at the cost of low throughput. For example, the RNASeqMetaDB provides a database of manually annotated terms associated with a set of mouse RNA-seq experiments (Zhengyu *et al*., 2015). This database describes only 306 RNA-seq experiments, which represents a small subset of all experiments in the SRA.

In contrast, automated annotation allows higher throughput at the cost of lower accuracy. Methods for automating the normalization of biomedical metadata frame the task as that of *entity recognition.* Entity recognition is the process of automatically recognizing and linking entities in natural language text to their corresponding entries in a controlled vocabulary. Tools that take this approach include ConceptMapper (Tanenblatt *et al*., 2010), SORTA (Pang *et al*., 2015), ZOOMA (Misha *et al*., 2012), and the BioPortal Annotator (Whetzel *et al*., 2011). Furthermore, there have been efforts to utilize such tools to automatically normalize large biomedical metadata sets. For example, work by Shah *et al.* (2009) automatically annotated samples and studies in the Gene Expression Omnibus (GEO) (Barrett *et al*., 2013) and other sources of biomedical metadata. Similarly, work by Galeota and Pelizzola (2016) annotated samples in GEO using ConceptMapper.

We assert that entity recognition alone is insufficient for automating the normalization of the SRA’s sample-specific metadata. Rather, since many of the sample’s descriptions mention ontology terms that describe extraneous entities (such as the study and experiment), a suitable solution should seek to extract *only* those terms that are being used to describe the biology of the sample. Biomedical entity recognition tools are best suited for data submitters who wish to facilitate annotation of their metadata before submission. Such tools do not adequately filter terms that do not describe the biology of the sample because they do not attempt to understand the fine-grained semantics of the text.

We further assert that important biological properties are often numerical and are not captured by ontology terms alone. Such terms include age, time point, and passage number for cell cultures. To the best of our knowledge, the problem of extracting real-value properties from metadata has yet to be addressed.

Lastly, we assert that ontology terms alone do not always provide enough context to understand the type of sample being described. For example, a cell culture that consists of stem cells differentiated into fibroblast cells may be annotated as both “stem cells” and as “fibroblast.” Such annotation leaves ambiguity as to whether the sample was differentiated from stem cells, or rather, was reprogrammed into a pluripotent state from primary fibroblasts. We assert that each sample should be categorized into a specific sample-type that captures the process that was used to obtain the sample.

To address these challenges, we present MetaSRA: a normalized encoding of biological samples in the SRA, along with the novel computational pipeline with which it was automatically constructed. MetaSRA encodes the metadata for each sample with a schema inspired by that used in the ENCODE project (Malladi *et al*., 2015). This schema is comprised of three parts:

1. Sample labels, using terms from the following biomedical ontologies: Disease Ontology (Kibbe *et al*., 2015), Cell Ontology (Bard *et al*., 2005), Uberon (Mungall *et al*., 2012), Experimental Factor Ontology (Malone *et al*., 2010), and the Cellosaurus (http://web.expasy.org/cellosaurus).
2. A sample-type classification, with six sample-type categories similar to those used by ENCODE.
3. Standardized numerical properties of the sample.

The first two parts are shared with the ENCODE schema, with the last part being a MetaSRA-specific extension. Currently, MetaSRA encodes all human samples utilized in RNA-seq experiments on the Illumina platform; however, future work will expand MetaSRA to other species and assays.

## 2 Data

We standardized all human samples assayed by RNA-seq experiments on the Illumina platform. An example of a standardized sample is shown in Figure 2. Metadata was retrieved from the SRAdb (Yuelin *et al*., 2013) downloaded on 09/05/2016. The BioSample’s sample-specific key-value pairs are stored in the “attribute” field of the “sample” table in the SRAdb. Our data set consists of 75038 samples, of which, 73407 are associated with a non-empty set of descriptive key-value pairs.

**Figure 2:**
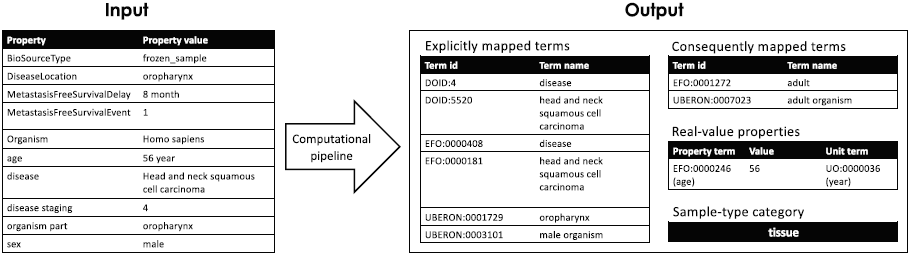
An example of the metadata normalization process for sample ERS183215. We extract explicit mappings, consequent mappings, real-value properties, and the sample-type category for each set of sample-specific key-value pairs in the SRA.

The samples processed belong to 2681 distinct studies and the number of samples contained in each study varies by several orders of magnitude (Figure 1C). These studies can be partitioned into 88 “large” (≥ 100 samples) and 2593 “small” (< 100 samples) studies, with the “large” studies constituting 57% of all samples processed. Due to the fact that samples belonging to a common study are described similarly, we argue that it is tractable to annotate samples belonging to the modest number of “large” studies using hand-tuned study-specific methods. In contrast, the large number of “small” studies and the diversity of their associated descriptions makes the process of designing study-specific methods for all of these studies intractable. We therefore focused our evaluations on samples belonging to “small” studies, with future work involving hand-tuning the normalization of samples from “large” studies.

## 3 Task definition

### 3.1 Mapping samples to ontologies

Like the ENCODE project, we label each biological sample using biomedical ontologies. An ontology is a structured knowledge-base that defines a set of concepts/terms within a specific domain of discourse. Besides providing the definition for each term, an ontology also encodes a directed graph in which each term is represented by a node and each edge represents a relationship between two terms. Edges are usually labelled with a relationship-type. For example, the most common edge is the `is_a` edge. Given terms *a* and *b*, *a* `is_a` *b* asserts that all instances of *a* are also instances of *b*. Similarly, the `part_of` edge represents the knowledge that one entity is a component of another entity. Labelling metadata using ontology terms allows for queries of the data that utilize the structured knowledge of the ontology. For example, a query for “brain” samples may return samples labelled with “cerebral cortex” because the cerebral cortex is a component of the brain.

We define the task of mapping samples to ontologies as follows: given a set of samples 𝓢, a set of ontology terms 𝓞, and set of relationship-types 𝓡, we seek a function ƒ: 𝓢 ⟶ 𝓟(𝓞), where 𝓟(𝓞) is the powerset of 𝓞, such that given a sample *s*, for each *ο* ∈ ƒ (*s*), there exists a relationship-type *r* ∈ 𝓡 that relates the sample *s* to the ontology term *ο*. We restrict 𝓡 to the following types of relationships:

- **has phenotype**: Maps samples to phenotypic or disease terms.
- **derives from**: Maps samples to cell lines or, when the sample consists of differentiated cells, to stem cell terms.
- **part of**: Maps samples to the anatomical entity from which it was extracted.
- **consists of cells of type**: Maps samples to their constituent cell types.
- **underwent**: Maps samples to ontology terms that describe a medical or experimental protocol.

We restrict our use of ontology terms to only “biologically significant” terms. An ontology term *o* is deemed biologically significant if given two samples *s*_1_, *s*_2_ ∈ *S* where ƒ(*s*_1_) ⊂ ƒ(*s*_2_) and ƒ(*s*_2_) \ ƒ (*s*_2_) = {*ο*} there likely exists a difference in gene expression or other measurable difference in biochemistry between the two samples. In simpler terms, an ontology term *ο* is deemed biologically significant if given two samples with equivalent descriptions barring that one sample can be described by *ο* and the other cannot, a significant difference in the biochemistry of the cell may exist between the two samples. For example, the ontology term for “cancer” is biologically significant, whereas the term “organism” is not because all samples are trivially derived from an organism. We map samples to only biologically significant terms in the ontologies.

#### 3.1.1 Discriminating between term mentions and term mappings

Our goal in mapping samples to ontology terms goes beyond named entity recognition. Rather than finding all occurrences or “mentions” of ontology terms in the metadata, we attempt to infer which labels adequately describe the biological sample being described. A term may be mentioned, but not mapped as well as mapped, but not mentioned.

For example, consider a sample’s description that includes the following text: `Metastatic castration resistant prostate cancer`. If we consider the Uberon and Disease Ontology, we see that the string “prostate cancer” mentions three terms in these ontologies: “prostate gland”, “cancer”, and “prostate cancer.” Of these terms, only “cancer” and “prostate cancer” are mapped because they are related to the sample through the “has phenotype” relationship. The string “prostate” is not not mapped because it localizes the disease rather than the sample. There is no relationship-type in 𝓡 that associates the sample with “prostate.” By prohibiting the mapping to “prostate”, we remove ambiguity as to whether the sample was derived from an organism with a prostate-related disease, or from prostate tissue itself. More generally, whenever a sample maps to an anatomical entity, we are asserting that the sample originated from that site.

To provide an example in which an ontology term should be mapped, but is not mentioned, consider a sample described by with the key-value pair `passage`: 4. The Cell Ontology term for “cultured cell” is not mentioned in this description; however, by the fact that it was explicitly stated that the cell was passaged, we can infer that the sample consists of cultured cells. Thus we map the sample to “cultured cell” via the “consists of cells of type” relationship.

#### 3.1.2 Discriminating between explicit and consequent mappings

We distinguish between two types of mappings: those that are explicit in the metadata and those that can be inferred from the explicit mappings. We refer to the latter as “consequent mappings.” For example, the ontology term for “female” is explicitly mapped from the key-value pair, `sex: female`, because the author is explicitly communicating the fact that this sample maps to “female” through the “has phenotype” relationship.

A sample “consequently” maps to an ontology term if, using external knowledge, one can logically conclude that the sample maps to the term. The premier example of such a case arises when a sample maps to a cell line. In such a case, the sample would consequently map to terms that describe this cell line. For example, given the key-value pair `cell line: MCF-7`, the sample would consequently map to “adenocarcinoma” because the MCF-7 cell line was established from a breast adenocarcinoma tumor. MetaSRA includes both explicit and consequent mappings.

### 3.2 Extracting real-value properties

In addition to mapping samples to ontology terms, we also annotate samples with real-value properties that are described in the metadata. We structure each real-value property as a triple (*property, value, unit*) where *property* is a property ontology term in the EFO, *value* ∈ ℝ, and *unit* is an ontology term in the Unit Ontology (Gkoutos *et al*., 2012). For example, the raw key-value pair age: 20 years old would map to the tuple (“age”, 20, “year”).

### 3.3 Predicting sample-type category

Like the ENCODE project, we categorize samples into their respective sample-type using categories similar to those used by ENCODE. These categories consist of `cell line, tissue, primary cell, stem cells`, and `in vitro differentiated stem cells, and induced pluripotent stem cell line`. Whereas ENCODE uses an `immortalized cell line` category, we instead use the category `cell line` to generalize to any cells that have been passaged multiple times, which include those from finite cell lines. Figure 5A illustrates how we define each sample-type category based on the methods by which the sample was obtained.

## 4 Methods

### 4.1 Mapping samples to ontologies

At the core of our method is a graph data structure for maintaining the provenance of each derived ontology term. This graph, which we call a Text Reasoning Graph (TRG), provides a framework for maintaining the provenance of extracted ontology terms, and for writing rules and operations that can reason about which terms should be mapped versus which are merely mentioned. Nodes in the TRG represent artifacts derived from the original metadata text. Such artifacts may be *n*-grams, inflectional variants, or synonyms. Other nodes in the graph represent mapping targets such as ontology terms or real-value property tuples. Edges between artifacts represent derivations from one artifact to another. An edge between an artifact and an ontology term represents a lexical match between the artifact and the ontology term. We implemented a computational pipeline that is composed of a series of stages that constructs the TRG. To start, the pipeline accepts the raw key-value pairs and constructs an initial TRG. Then, each stage operates on the TRG by modifying its nodes and edges. Figure 3 depicts the subgraph of a final TRG that maps a key-value pair to a set of ontology terms. By maintaining the provenance of each derived ontology term we can implement custom reasoning operations that more accurately determine which terms describe the sample. Such reasoning operations utilize the graph structure to filter out ontology terms for which there is no relationship-type in 𝓡 that describes the relationship between the sample and the ontology term.

**Figure 3:**
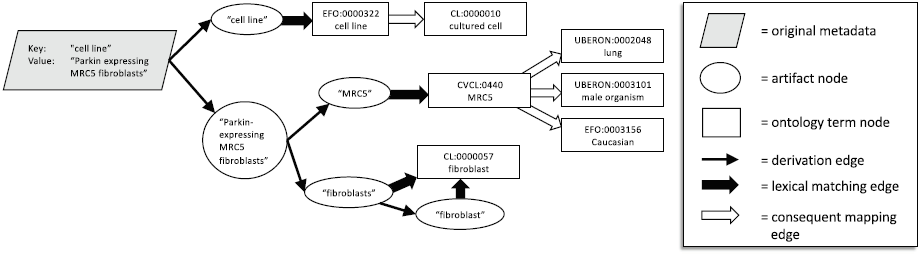
A subgraph of the TRG constructed from sample SRS1212219 illustrating the graph data structure that our pipeline maintains as it reasons about the sample. This framework allows us to maintain the context of each artifact. For example, we map to the MRC5 cell line only because there is a mapping to the “cell line” ontology term in the graph emanating from the key. We also note the terms for “lung”, “male organism”, and “Caucasian” were mapped to the MRC5 cell line from the ATCC cell bank data and are thus *consequent mappings*.

In the following sections we describe the most notable stages of the pipeline. Full details are provided in the supplementary materials.

#### 4.1.1 Filtering key-value pairs

Before initializing the TRG, we filter key-value pairs from the metadata where either the key or value appears in a set of blacklisted keys and values. This blacklist of keys contains those that describe a property that does not pertain to the biology of the sample, such as “study name” and “biomaterial provider.” The blacklist of values include those that negate the key, such as “none” or “no.” For example, the key-value pair `is tumor: no` is removed because the value `no` negates the property `is tumor`.

#### 4.1.2 Artifact generation

We define an artifact to be any string that is derived from a substring of the original metadata text. Such artifacts include include n-grams, lower-cased words, and inflectional and spelling variants of words in the metadata. An artifact node in the TRG represents a single artifact. Several stages of the pipeline generate new artifact nodes from existing artifact nodes and draw edges from original to derived artifacts. One such stage derives inflectional and spelling variants from existing artifacts using the National Library of Medicine’s SPECIALIST lexicon (Browne *et al*., 2000). For example, given an artifact node representing the pluralized noun “fibroblasts”, this stage will create a node for the singular noun “fibroblast” and draw a directed edge from “fibroblasts” to “fibroblast.”

#### 4.1.3 Matching artifacts to ontologies

We perform fuzzy string matching between all artifacts and the ontology terms to find all exact matches and minor misspellings (for misspelling criteria, see supplementary materials). In general, fuzzy matching is a computationally expensive task. To speed up this process, we pre-compute a metric tree index for all ontology term names and synonyms as described by Bartolini *et al.* (2002). The index allows us to filter for ontology term strings that are nearby the query string in edit space. We then explicitly compute the edit distances to these nearby strings.

#### 4.1.4 Graph reasoning

Certain stages of the pipeline utilize the structure of the TRG. We refer to such steps as “reasoning” steps. For example, we remove extraneous mappings to cell line terms by searching the graph emanating from the key for a lexical match to ontology terms such as “cell line” and “cell type.” If such a match is *not* found, we search the graph emanating from the value for artifacts that have a lexical match to a cell line ontology term and remove all such ontology term nodes. This process is important for removing false positives due to the fact that names of cell lines are often similar to gene names and acronyms. For example, “Myelodysplastic Syndromes” is often shortened to “MDS.” MDS also happens to be a cell line in the Cellosaurus.

To provide another example, we create a list of “context-specific synonyms” and derive synonyms for artifacts only when the provenance of the artifact meets a certain criterion. For example, a common key-value pair is `sex: F`. Here, the string “F” is an abbreviation for “female”; however, this is only known because the key describes the sex of the organism. “F” in another context may not be an abbreviation for “female.” One stage of our pipeline derives synonyms for artifacts only when the given artifact was extracted from a key-value pair describing a specific property.

#### 4.1.5 Mapping cell line terms using ATCC cell bank data

Our pipeline draws edges between a cell line ontology term node and the ontology terms that describe the cell line. For example, if the TRG contains the node for the cell line “HeLa”, we draw an edge to the ontology terms for “adenocarcinoma” and “female” because this cell line was derived from a woman with cervical adenocarcinoma. We consider such mappings to be consequent mappings because they are retrieved using an external knowledge base. In this case, the external knowledge base was created from data we scraped from the ATCC website (https://www.atcc.org). To construct mappings between cell lines and ontology terms, we ran a variant of our pipeline on the scraped cell line data. We scraped cell line metadata for all cell lines that are present in the Cellosaurus.

#### 4.1.6 Maximal phrase-length mapping

It is a common occurrence for disease ontology terms to include anatomical entities in their name. For example, “breast cancer” includes “breast” as a substring. As previously discussed, under our framework, it would be incorrect to map “breast” to a sample solely based on a mention of “breast cancer” in its metadata because “breast” localizes the cancer, but does not localize the origin of the sample. Whereas it is entirely possible that such a sample was indeed derived from breast tissue, without additional information we cannot eliminate the possibility that the sample originated from some other tissue, such as from a malignant site. In this example, we maintain a conservative approach and avoid mapping to “breast.” We implement this process by having each artifact node keep track of the original character indices in the metadata from which it was derived. After mapping all artifacts to the ontologies, we remove all ontology terms that were lexically matched with an artifact node that is subsumed by another artifact node that matches with an ontology term.

#### 4.1.7 Linking ontologies

The domain covered by the EFO overlaps with many of the other ontologies because it includes cell types, anatomical entities, diseases, and cell lines. In many cases, the EFO is inconsistent with other ontologies in how it draws edges between terms. For example, the term “lung adenocarcinoma” and “adenocarcinoma” are present in both the Disease Ontology and the EFO; however “adenocarcinoma” is a parent of “lung adenocarcinoma” in the Disease Ontology but not in the EFO. These inconsistencies pose a problem when we apply our maximal phrase-length mapping process. For example, when a sample maps to “lung adenocarcinoma” and “adenocarcinoma”, we remove “adenocarcinoma” because it is a substring of “lung adenocarcinoma.” This is valid for the Disease Ontology because the term for “adenocarcinoma” is implied by “lung adenocarcinoma” by its position in the ontology. However, this results in a false negative for the EFO version of this term.

To counteract this problem, we link EFO terms to terms in the other ontologies. Two terms are linked when they share the same term-name or exact-synonym. Then, when an artifact maps to a term, we traverse the term’s ancestors and map to any terms that are linked to those ancestors. In the case of “lung adenocarcinoma”, we would traverse the ancestors of this term in the Disease Ontology and map to the EFO’s “adenocarcinoma” because it is linked to the Disease Ontology version of this term.

### 4.2 Extracting real-value properties

We maintain a list of ontology terms that define real-value properties. Currently, we use 7 terms including “age”, “passage number”, and “time-point.” Future work will entail expanding this list. To extract a real-value property from a key-value pair, we search the graph emanating from the key for a match to a property ontology term. If such a property is found, we search the graph emanating from the value for an artifact that represents a numerical string as well as a unit ontology term node (for example “46” and “year”). From this process, we extract the triple (property, value, unit). For example, given the key-value pair `age: 46 years old`, we extract (“age”, 46, “year”).

### 4.3 Predicting sample-type category

We classify samples by sample-type using a supervised machine learning approach. We trained a one-vs.-rest ensemble of logistic regression, binary classifiers where each classifier was trained using L1 regularization.

#### 4.3.1 Training set

We manually annotated 705 samples based on their metadata. We determined each sample’s sample type by consulting the sample’s study, publication, and other external resources that describe the experimental procedure used to obtain the sample. Since samples that belong to the same study are likely described similarly, one potential pitfall in the learning process is that if training samples are drawn uniformly and at random from all samples, the learner will be biased towards features that correlate with how larger studies describe their samples rather than features that correlate with sample-type. To avoid this issue, we ensured that no two samples in the training set came from the same study.

#### 4.3.2 Feature selection

We consider two types of features for representing each sample: *n*-gram features and ontology term features. For *n*-gram features, we consider all uni-grams and bi-grams appearing in the training samples’ raw metadata. For ontology term features, we consider the set of all ontology terms that were mapped to the training samples by our automated pipeline. We performed a feature selection process (for details, see supplementary materials) that involved using mutual information to select features that are indicative of at least one of the target sample-types.

#### 4.3.3 Prediction

When making a prediction on an unseen sample *x*, each binary classifier *c_j_* in the ensemble computes its estimate of the conditional probability *c_j_* (*x*):= *p*(*y* = *j*∣*x*), which can be interpreted as the confidence that classifier *c_j_* believes that *x* is of type *j*. Using these probabilities, we designed a decision procedure that utilizes our domain knowledge for determining the sample type. We found that injecting domain knowledge into the process boosted performance in cross-validation experiments. This decision procedure limits the possible sample-types based on the ontology terms mapped to the sample. For example, if “stem cell” was mapped to the sample, we limit the possible predictions to `stem cells, induced pluripotent stem cell line`, and `in vitro differentiated cells`. Although theoretically, the learning algorithm should learn these facts itself, there is likely not enough training examples for the algorithm to learn such patterns.

## 5 Results

### 5.1 Evaluation of ontology mappings

In order to create a test set for evaluation of our pipeline, we manually normalized metadata for 206 samples from the SRA where each sample belongs to a unique study. These samples were recently added to the archive and had not been considered during the development of our computational pipeline. Thus, performance on this subset of data provides an unbiased estimate of its ability to generalize to unseen samples.

#### 5.1.1 Evaluating explicitly mapped ontology terms

We first evaluated our pipeline’s ability to map samples to explicitly mapped ontology terms using the following metrics: recall, error rate, specific terms recall, and specific terms error rate. Given a sample, let *T* be the set of all ontology terms to which the sample maps including terms ancestral to those explicitly mentioned. Let *T*' be the most specific terms in *T*. That is *T*':= {*t* ∈ *T*: no child of *t* is in T}. Let *P* be the set of predicted terms to which the sample maps. Let *P*' be the most specific terms in *P*. That is *P*':= {*p* ∈ *P*: no child of *p* is in P}. We define our metrics as follows:

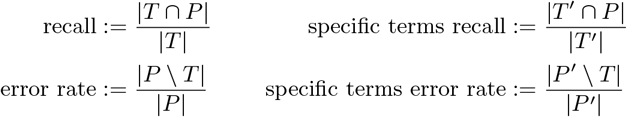

We use these four metrics instead of traditional precision and recall because precision and recall are affected by the structure of the ontology. This is due to the fact that the act of retrieving an ontology term implicitly retrieves all of its ancestral terms in the ontology’s directed acyclic graph. Thus, retrieving a term with a high number of ancestral terms will lead to exaggerated metrics. The specific terms error rate corrects for this by describing the fraction of the most specific predicted terms that incorrectly describe the sample. We note that the error rate is simply 1 – precision. The metrics are demonstrated in Figure 4A.

**Figure 4:**
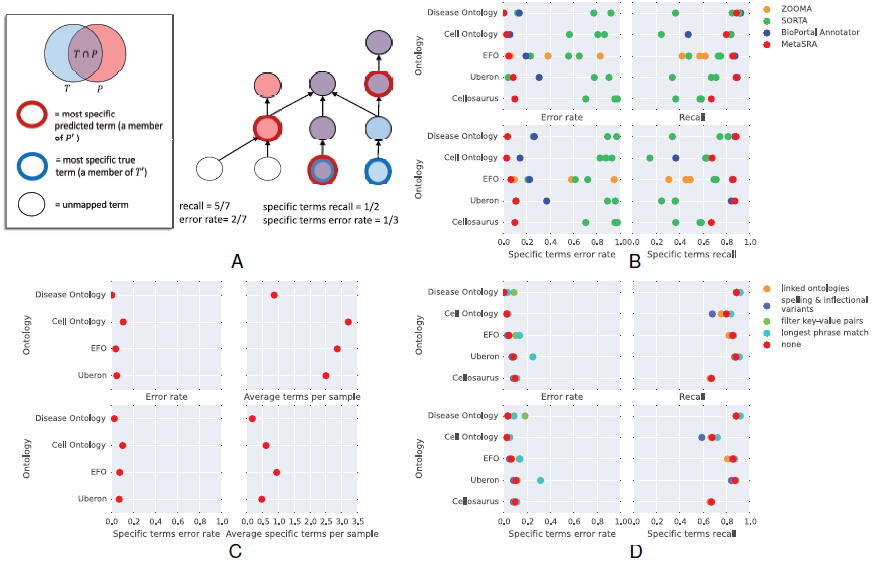
(A) A schematic of an ontology subgraph demonstrating our calculation of recall, specific terms recall, error rate, and specific terms error rate. (B) Recall, error rate, specific terms recall, and specific terms error rate for versions of our pipeline in which certain stages are disabled. The data points labelled “none” refer to the complete pipeline in which no stage is disabled. (C) The error rate, specific terms error rate, average retrieved terms per sample, and average specific retrieved terms per sample across all ontologies when considering only consequently mapped terms. (D) Performance of our pipeline in mapping explicit ontology terms versus BioPortal’s Annotator, ZOOMA, and SORTA. We measured recall, error rate, specific terms recall, and specific terms error rate for all programs across all ontologies with the exceptions that ZOOMA only maps to the EFO and only MetaSRA and SORTA map to the Cellosaurus.

We compared our pipeline to the BioPortal Annotator, SORTA, and ZOOMA. We ran SORTA using the three confidence thresholds of 1.0, 0.5, and 0.0. We also ran ZOOMA using the three confidence thresholds of high, good, and low. The metrics across all programs and ontologies are shown in Figure 4B.

Our pipeline outperformed other programs in specific terms error, and specific terms recall across all ontologies. The pipeline’s recall was similar to that achieved by the BioPortal Annotator. However, since the BioPortal Annotator simply detects ontology mentions, many of these mentions are false positives because they do not describe the sample.

#### 5.1.2 Evaluating the impact of individual pipeline stages

In order to evaluate the performance impact of each of our pipeline’s stages, we ran our pipeline on our test set with certain individual stages disabled. Figure 4C shows the performance impact when removing stages for filtering key-value pairs, linking ontologies, filtering sub-phrase matches, and generating spelling and inflectional variants. As expected, the results of these tests indicate a general trade off between recall and error rate. Certain stages may decrease recall, but pose the benefit of decreasing the error rate. Furthermore, a given stage may be more effective for mapping terms in some ontologies rather than others.

#### 5.1.3 Evaluating consequently mapped ontology terms

Recall is an inappropriate metric for evaluating our ability to map consequent terms due to the fact that the set of consequent terms is undefined. By our definition, a consequent term is any term that the sample can be mapped to based on expert or external knowledge. Thus, depending on the expert or external knowledge base, the set of consequent terms may change. Furthermore, an expert may use an exceedingly large number of ontology terms to describe the sample depending on what she knows about the sample and experiment. For these reasons, we look at the total average number of consequent terms that we map to each sample. This metric describes the amount of extra information that is provided when considering external knowledge. We further looked at the average number of most specific mapped consequent terms. Figure 4D displays these metrics across all ontologies. We note that no terms from the Cellosaurus were consequently mapped and are thus not displayed in the figure.

### 5.2 Evaluating extraction of real-value properties

Of the 206 samples in our test set, 63 described real-value properties. We use precision and recall to evaluate our performance in retrieving real-value property tuples. We call a predicted real-value property a true positive if the property type, value, and unit all match the ground truth. On these 62 samples, we report precision of 1.0 and recall of 0.487.

### 5.3 Evaluating sample-type predictions

To evaluate our ability to predict each sample’s sample-type, we evaluated the algorithm’s performance on a held-out test set. This test set consists of the same samples that were utilized for evaluating our ontology term mapping procedure. We annotated all samples for which the origin of the sample was explained in an external resource such as a scientific publication. In total, this came to 178 samples where no two samples belong to the same study. The distribution of sample-types in our test set is illustrated in the bar graph above the matrix in Figure 5B.

**Figure 5:**
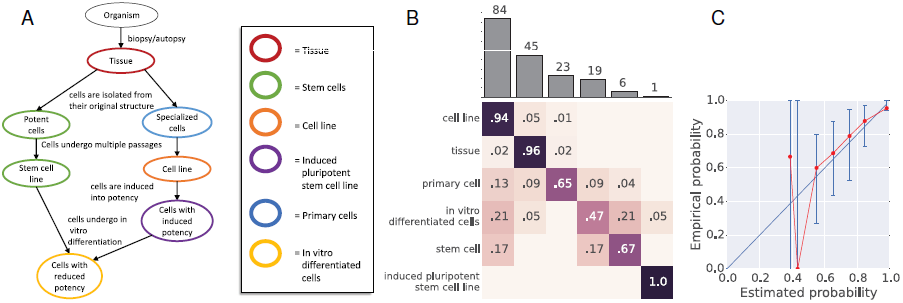
(A) A graph illustrating how sample-type categories are defined. Each node in the graph represents a biological sample. Arrows between nodes represent procedures carried out on the sample. Nodes are colored according to their sample-type category. (B) Confusion matrix for sample-type category prediction accuracy on the test data set. Above the confusion matrix we display the number of samples in each category. (C) Calibration of the model. The estimated probability of the model (average of confidence values in each bin) is plotted against the empirical probability that the model is correct (accuracy of predictions in each bin). The straight blue-line plots a well-calibrated model. Error bars are drawn according to a bootstrap sampling approach described in Bröcker and Smith (2007). No points are drawn for bins that contain no predictions.

Our trained classifier achieved an accuracy of 0.848 over these samples. The confusion between between categories is plotted in the confusion matrix in Figure 5B. In general, the classifier does well in determining the cell line samples and tissue samples. Close inspection of the classifier’s errors revealed that most were due to samples with poor quality descriptions. Such samples are difficult to categorize, even as a human, without consulting the scientific publication in which the sample is described. Correctly classifying these samples will require utilizing external descriptions of the samples.

MetaSRA includes the classifier’s confidence of each prediction. To evaluate the quality of these confidence scores, we assessed the calibration of the classifier. A classifier is well calibrated if for any instance *x* with true label *y*, it holds that 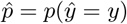 where 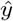 is the classifier’s predicted class label of *x* and 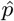 is its confidence. To assess calibration, we group the predictions into bins according their confidence scores. In Figure 5C, we plot the average confidence of predictions in each bin (i.e. estimated probability) against the empirical accuracy (i.e. empirical probability) of predictions in that bin. A well calibrated model should produce points near the line with slope 1 and y-intercept 0. Error bars were constructing using a bootstrapping method described by Bröcker and Smith (2007).

## 6 Availability

MetaSRA is available at http://deweylab.biostat.wisc.edu/metasra in both an SQLite database as well as in a JSON text file. Each update to MetaSRA will be accompanied with a unique version number that can be used to identify each discrete version of the normalized metadata. The code used for creating MetaSRA has been open-sourced at https://github.com/deweylab/metasra-pipeline.

## 7 Discussion and future work

Although previous work has addressed the task of annotating biomedical text, there had yet to be a thorough effort at generating an accurate annotation of sample-specific metadata for the SRA. MetaSRA addresses this gap, providing normalized metadata encoded into a schema inspired by that used by the ENCODE project. Currently, the MetaSRA includes normalized sample-specific metadata for human samples assayed by RNA-seq experiments on the Illumina platform. We expect that this resource will enable higher utilization of the SRA and investigations across diverse phenotypes, diseases, cell types, and conditions.

Future work will involve expanding MetaSRA to incorporate more biological samples as well as to expand the set of ontologies used for annotation. First, we plan to expand MetaSRA to all human samples (not only those used in RNA-seq experiments) as well as to samples from other species. We will also expand the set of ontologies to those that include experimental variables. For example, we hope to include the ChEBI ontology for annotating samples with chemical treatments (Hastings *et al*., 2013). Our initial experiments with ChEBI revealed that mapping samples to this ontology poses unique challenges because of its relatively large size and inclusion of synonymous terms that share names with those representing unrelated concepts (e.g., the ChEBI term “maleate(2-)” has the synonym “male”). We also plan to incorporate more sources of external knowledge in the MetaSRA construction pipeline in order to map samples to a larger number of consequent terms. For example, additional sources of cell line data, such as the Coriell Institue for Medical Research (https://www.coriell.org), could be used to annotate cell lines that are not present at the ATCC. Lastly, we plan to leverage other sources that provide information on *sets* of samples, such as a publication describing the study in which a set of samples was assayed. Although such sources provide valuable details, because they are not sample-specific, a key challenge will be to automatically determine which details may be confidently assigned to individual samples.

